# Temporal Genetic Dynamics of an Experimental, Biparental Field Population of *Phytophthora capsici*

**DOI:** 10.1101/089953

**Authors:** Maryn O. Carlson, Elodie Gazave, Michael A. Gore, Christine D. Smart

## Abstract

Defining the contributions of dispersal, reproductive mode, and mating system to the population structure of a pathogenic organism is essential to estimating its evolutionary potential. After introduction of the devastating plant pathogen, *Phytophthora capsici*, into a grower’s field, a lack of aerial spore dispersal restricts migration. Once established, coexistence of both mating types results in formation of overwintering recombinant oospores, engendering persistent pathogen populations. To mimic these conditions, in 2008, we inoculated a field with two *P. capsici* isolates of opposite mating type. We analyzed pathogenic isolates collected in 2009-13 from this experimental population, using genome-wide single-nucleotide polymorphism markers. By tracking heterozygosity across years, we show that the population underwent a generational shift; transitioning from exclusively F_1_ in 2009-10; mixed generational in 2011; and ultimately all inbred in 2012-13. Survival of F_1_ oospores, characterized by heterozygosity excess, coupled with a low rate of selfing, delayed declines in heterozygosity due to inbreeding and attainment of equilibrium genotypic frequencies. Large allele and haplotype frequency changes in specific genomic regions accompanied the generational shift, representing putative signatures of selection. Finally, we identified an approximately 1.6 Mb region associated with mating type determination, constituting the first detailed genomic analysis of a mating type region (MTR) in *Phytophthora*. Segregation patterns in the MTR exhibited tropes of sex-linkage, where maintenance of allele frequency differences between isolates of opposite mating type was associated with elevated heterozygosity despite inbreeding. Characterizing the trajectory of this experimental system provides key insights into the processes driving persistent, sexual pathogen populations.

## Introduction

*Phytophthora capsici* is the filamentous, soil-borne oomycete plant pathogen responsible for Phytophthora blight, a disease inflicting significant annual crops losses worldwide (Erwin and Ribeiro, 1996; Granke et al., 2012; Hausbeck and Lamour, 2004; Lamour et al., 2012). Success of *P. capsici* is facilitated by its widespread ability to overcome fungicides (Lamour and Hausbeck, 2000), dearth of resistant cultivars (Granke et al., 2012), and large, diverse host range (comprising >15 plant families), including widely grown, economically important vegetable crops in the *Cucurbitaceae, Solanaceae,* and *Fabaceae* plant families (Hausbeck and Lamour, 2004; Satour and Butler, 1967; Tian and Babadoost, 2004). Extreme weather events often initiate new infestations by introducing inoculum into agricultural fields via flood waters (Dunn et al., 2010). Contaminated soil and infected plant material are commonly implicated in pathogen spread (Granke et al., 2012), however, *P. capsici* is not aerially dispersed (Granke et al., 2009).

Once introduced into a field, the explosive asexual cycle of *P. capsici* catalyzes the rapid escalation of disease within a growing season. When exposed to water saturated conditions, a single sporangium can release 20-40 zoospores, each capable of inciting root, crown, or fruit rot, the characteristic symptoms of Phytophthora blight (Hausbeck and Lamour, 2004). For sexual reproduction, the heterothallic *P. capsici* requires two mating types, classically referred to as A1 and A2 (Erwin and Ribeiro, 1996). Exposure to mating type specific hormones (α1 and α2) stimulates production of the gametangia, subsequent outcrossing, and formation of recombinant oospores (Ko, 1988). However, both mating types produce both male and female gametangia, and thus are capable of self-fertilization (Ko, 1988; Shattock, 1986), which is thought to occur at a lower rate relative to outcrossing in *P. capsici* (Dunn et al., 2014; Uchida and Aragaki, 1980).

While the asexual reproductive cycle directly inflicts crop damage, sexual reproduction confers several epidemiological advantages. First, unlike asexual propagules, oospores survive exposure to cold temperatures (Babadoost and Pavon, 2013; Hausbeck and Lamour, 2004). Thus, in regions with cold winter conditions, oospores are the primary source of overwintering inoculum (Bowers, 1990; Granke et al., 2012; Lamour and Hausbeck, 2003). Second, oospores remain in the soil for years regardless of host availability, enabling the persistence of the pathogen between growing seasons and rendering eradication unfeasible. In the spring, in the presence of susceptible hosts, germinating oospores, potentially formed in distinct years, initiate the repeating, asexual reproductive cycle (Granke et al., 2012; Hausbeck and Lamour, 2004).

Where both mating types coexist, sexual reproduction is associated with persistent pathogen populations, genetic diversity, and an approximate 1:1 ratio of A1 to A2 mating types (Dunn et al., 2010; Lamour and Hausbeck, 2001). While asexual reproduction can increase the prevalence of a specific genotype within a sexually reproducing population, the inability of asexual propagules to survive cold winters (Babadoost and Pavon, 2013; Hausbeck and Lamour, 2004) implies that each year meiosis disrupts linkage between the particular combination of alleles observed within a clone (Kondrashov, 1988). As a consequence, sexual reproduction mediates the effects of clonal propagation on *P. capsici* population structure (Lamour and Hausbeck, 2001). Furthermore, in geographic regions where sexual reproduction occurs, genetic differentiation between field populations, even within close proximity, suggests that after an initial introduction limited gene flow occurs between fields (Dunn et al., 2010; Lamour and Hausbeck, 2001), consistent with a lack of aerial dispersal (Granke et al., 2009).

Given this infection scenario, i.e. an initial inoculation but no subsequent introductions, we would expect *P. capsici* populations to exhibit signatures of a bottleneck event: reductions in genetic diversity and an increase in inbreeding over time, proportional to the number of founding isolates (Kirkpatrick and Jarne, 2000). (We define inbreeding strictly as inter-mating between related isolates, and reserve selfing to refer to self-fertilization events.) In populations which undergo a so-called founder effect, inbreeding is expected to decrease mean population fitness over time due to the expression of recessive deleterious alleles, i.e. the genetic load, in the homozygous state (Charlesworth and Charlesworth, 1987; Hartl and Clark, 2007). A related phenomenon, inbreeding depression, i.e. the difference in fitness between selfed and outcrossed progeny in a population (Kirkpatrick and Jarne, 2000), is considered a major driver of obligate outcrossing, and may contribute to maintenance of self-incompatibility in hermaphroditic plant species (Charlesworth and Charlesworth, 1987). Charting the genetic trajectory of isolated populations of *P. capsici* in the context of these processes, is essential to understanding pathogen evolution in an agriculturally relevant scenario.

Thus, in 2008, to investigate the response of *P. capsici* to a severe bottleneck, we established a closed, biparental field population, by inoculating a research field once with two heterozygous strains of opposite mating types. In a preliminary study, we tracked the allele and genotypic frequencies of five microsatellite markers in the field population from 2009-12 (Dunn et al., 2014). We demonstrated that sexual reproduction resulted in high genotypic diversity, a function of the proportion of unique isolates (Grünwald et al., 2003), in 2009-11, with a reduction in genotypic diversity in 2012. However, five markers afforded limited statistical power to characterize population and individual level phenomena.

Therefore, in the present study, we analyzed isolates collected in 2009-13 from the *P. capsici* field population with genotyping-by-sequencing (GBS), a multiplexed reduced-representation sequencing technique, which simultaneously identifies and scores single nucleotide polymorphism (SNP) markers distributed throughout the genome (Elshire et al., 2011). The closed experimental field design excluded introduction of new alleles via migration, providing a unique opportunity to address the influence of inbreeding on population genetic phenomena in *P. capsici.* In high-density SNP genotyping isolates from the biparental field population, our goal was threefold: 1) Evaluate the effects of oospore survival on population structure; 2) Quantify the genome-wide incidence of inbreeding; and 3) Identify whether specific regions deviate from the rest of the genome in terms of changes in allele frequency.

## Results

### GBS of the experimental biparental isolates and validation

We genotyped 232 isolates collected from a closed, biparental field population of *P. capsici* from 2009-13. All field isolates were collected from infected plant tissue, and are therefore, by definition, pathogenic. Additionally, we genotyped 46 single-oospore progeny from an *in vitro* cross between the same founding parents as a reference for the field isolates, for which generation was *a priori* unknown. Three of the *in vitro* progeny were identified as putative selfs by Dunn et al. (2014), which was confirmed by our analysis (see ‘Selfing in the lab and field’), and are hereafter referred to as *in vitro* selfs to distinguish them from the *in vitro* F_1_ progeny. The A1 (isolate: 0664-1) and A2 (isolate: 06180-4) founding parents were genotyped 14 and 11 times, respectively, to estimate laboratory and genotyping errors (Supplementary table S1).

Out of the 401,035 unfiltered variant calls, initial site filters reduced the data set to 23,485 high-quality SNPs (Supplementary Figure S1), with an average SNP call rate (i.e. the percentage of individuals successfully genotyped at each SNP) of 95.93% (median of 97.64%). The 23,485 SNPs were equally distributed among 307 scaffolds (scaffold size and number of SNPs were highly correlated (*r*^2^=0.95)), with an average SNP density of approximately 1 SNP every 2.5 kb. There was essentially no correlation between mean individual read depth and heterozygosity per SNP among all isolates (*r*^2^=0.009, *P*-value=0.10), indicating that heterozygous calling post-filtering was robust to differences in mean individual sequencing coverage (Supplementary Figure S2). Genotype files are available in VCF format upon request.

To assess SNP genotyping accuracy, we compared biological and technical replicates of the parental isolates. Replicates of the A1 parent (*n*=14) and A2 parent (*n*=11), representing 4-5 distinct serial cultures, shared on average 98.30% (*s*=0.45%) and 98.17% (*s*=0.51%) alleles identity-by-state (IBS), respectively (Supplementary Table S1). This corresponded to 3.60% (*s*=0.86%) discordant sites on average among non-missing genotypes between replicates. Lower average discordance 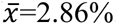, *s*=0.33%) between only replicates of the same parental culture (*n*=54 pairwise comparisons) suggested variation associated with distinct culture time points. Therefore, our overall genotyping error rate, inclusive of variation in mycelial and DNA extractions, but not different culture time points, was approximately 3%. Among technical replicates [same DNA sample (*n*=4) sequenced 3-4 times] the error rate was on average 2.95%, indicating that most of the genotype discrepancies were attributed to sequencing and genotyping errors rather than distinct mycelial harvests. When we excluded heterozygous calls in each pairwise comparison (*n*=21) of the technical replicates, less than 0.0001% sites were discordant, indicating that heterozygote genotype discrepancies drove genotyping errors. As in the total data set, the association between individual sequencing coverage and heterozygosity was negligible in both sets of parental replicates (Supplementary Figure S2).

*Phytophthora capsici* reproduces asexually, therefore, it was theoretically possible to sample the same genotype from the field multiple times within a year. To remove the bias imparted on population genetic analyses by including clones, we retained only one isolate for each identified unique genotype (Milgroom, 1996). Pairwise identity-by-state (IBS) between replicates of the A1 and A2 parental isolates were compared to establish a maximum genetic similarity threshold to define clones (see Methods), akin to (Meirmans and van Tienderen, 2004; Rogstad et al., 2002). Applying this threshold, we identified 160 unique field isolates out of the initial 232 field isolates (Supplementary Table S2). Two *in vitro* isolates and one field isolate were identified as outliers with respect to deviation from the expected 1:1 ratio of allele depths at heterozygous sites (*n*=2) or heterozygosity (*n*=1), and subsequently removed (Supplementary Figure S3). Previous studies have shown that deviation from a 1:1 ratio of allele depths at heterozygous sites, the expectation for diploid individuals, is correlated with ploidy variation (Li et al., 2015; Rosenblum et al., 2013; Yoshida et al., 2013), therefore the two allele depth ratio outliers provide preliminary evidence for ploidy variation in *P. capsici.* After outlier removal, the final data set consisted of 159 field isolates, 41 *in vitro* F_1_, and three *in vitro* selfs.

Clones did not appear in multiple years, consistent with the inability of asexual propagules to survive the winter (Babadoost and Pavon, 2013; Hausbeck and Lamour, 2004). After clone-correction, the A2 mating type was more represented in the field (A1:A2=65:94; *χ*2 test, *P*-value=0.02), a phenomenon also observed in the *in vitro* F_1_ (A1:A2=16:25; %2 test, *P*-value=0.16; Figure 1). The only exception was 2012, which may be explained by a smaller sample size in this year, artificially compounded by loss of several unique isolates (based on microsatellite profiles (Dunn et al., 2014) in culture prior to this study.) We observed lower genotypic diversity in 2012-13 (Supplementary Table S2), consistent with Dunn et al. (2014).

**Figure 1.**
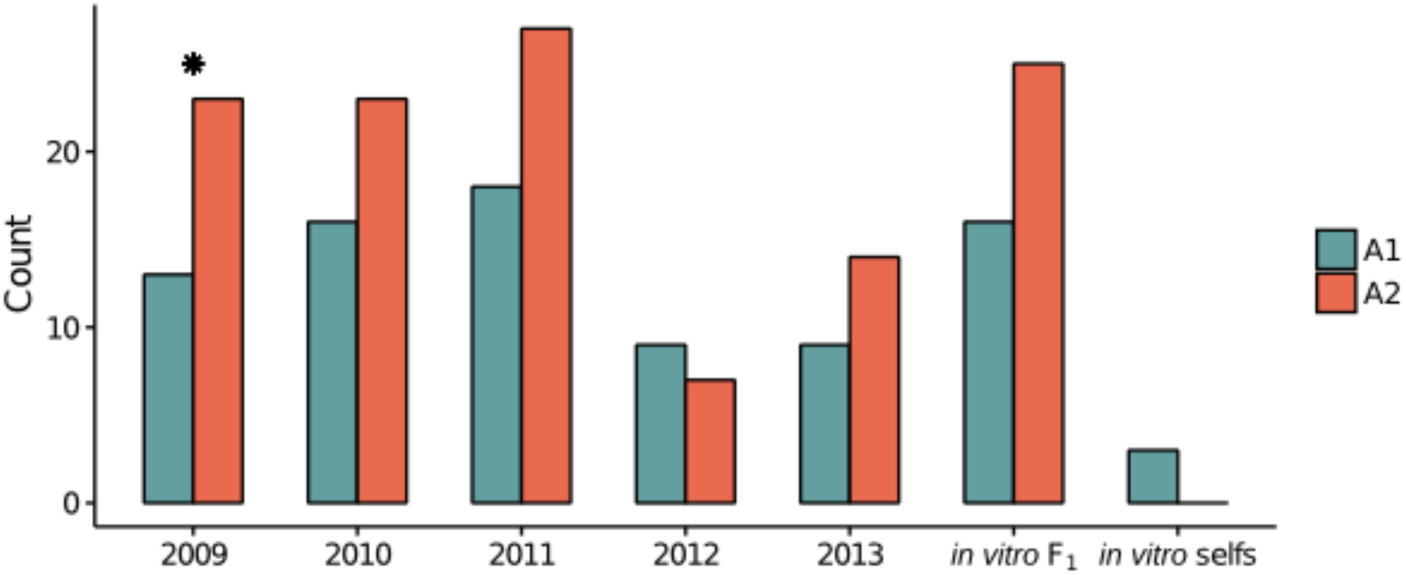
Distribution of the mating type of each isolate by year in the final, clone-corrected data set. Counts of the mating type of each isolate, A1 (teal) and A2 (reddish brown), in the *in vitro* F_1_, *in vitro* selfs, and clone-corrected field isolates, separated by year. The star indicates a significant difference (*χ*2 test; *P*-value<0.1) between Al and A2 counts.

To reduce oversampling of specific genomic regions, which can disproportionately influence population genetic inference (Abdellaoui et al., 2013; Price et al., 2006), without making assumptions about linkage disequilibrium (LD), we randomly selected one SNP within a given, non-overlapping 1 kb window. With final quality filters, and including only SNPs in scaffolds containing at least 300 kb (*n*=63), pruning resulted in a data set of 6,916 SNPs (Supplementary Figure S1). Bimodal heterozygosity and minor allele frequency (MAF) distributions in this reduced SNP set were consistent with distributions in the unpruned data set (Supplementary Figure S4). The pruned data set had a median SNP call rate of 98.01% and median site depth of 18.61 (i.e. average number of reads per individual per SNP). The median sample call rate (i.e. percentage of SNPs genotyped in each sample) was 97.77%, and the median sample depth (i.e. average number of reads per SNP per individual) was 20.36. Among technical replicates (*n*=4) the error rate was on average 1.52%. We utilized the pruned data set for all subsequent analyses.

### Population differentiation increases with year

To broadly define genetic relationships between the *in vitro* and field isolates relative to the founding parents, we analyzed the field, *in vitro* and parents (represented by consensus parental genotypes, see Methods) jointly, with principal component analysis (PCA). The PCA exhibited the expected biparental population structure, in that the majority of isolates clustered in between the parental isolates along the major axis of variation, principal component (PC) 1 (Figure 2A). Most 2009-11 isolates clustered with the *in vitro* F_1_, whereas, many 2012-13 isolates were dispersed along both axes, suggesting differentiation associated with year.

**Figure 2.**
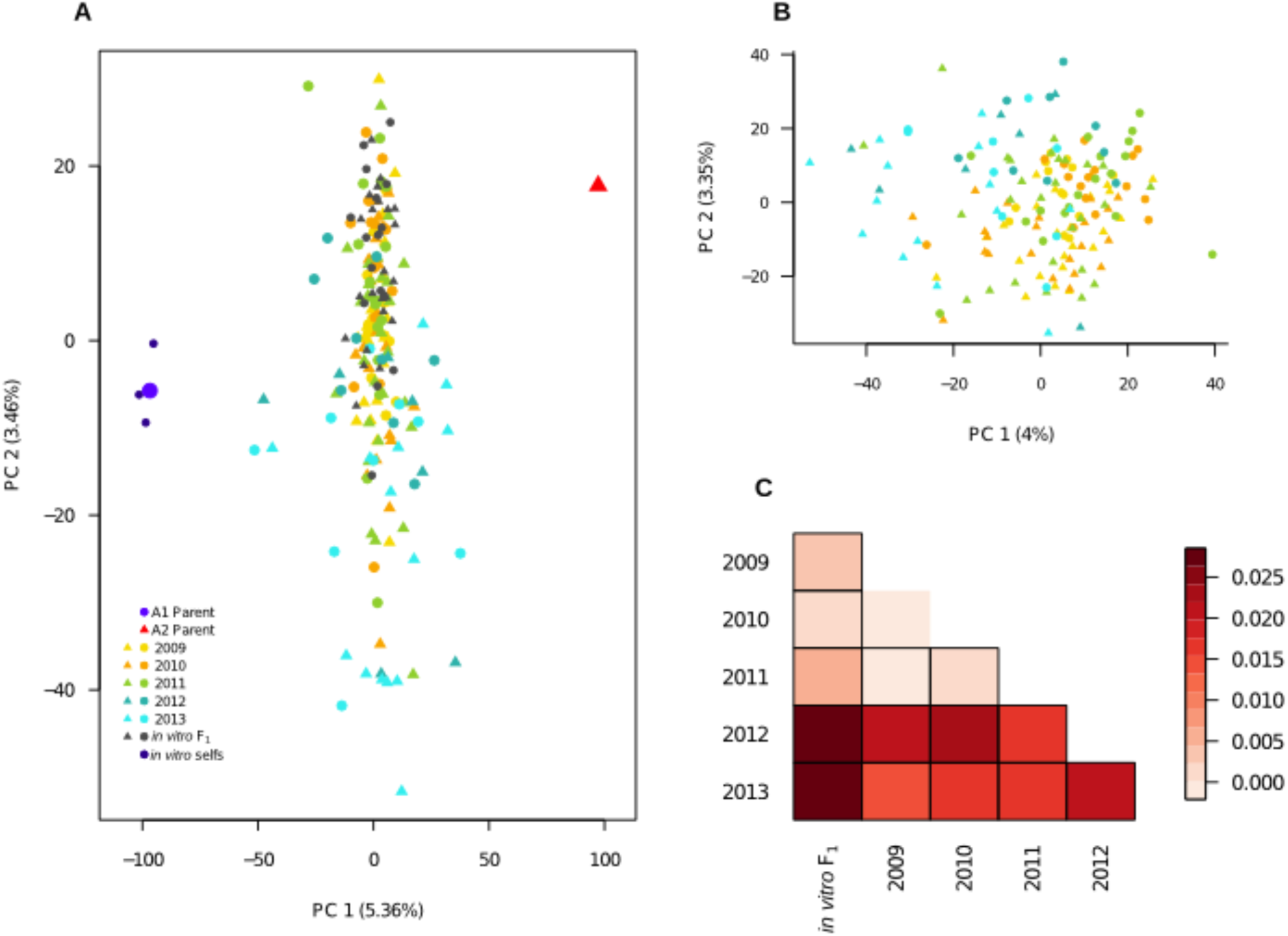
Population structure in the biparental field population relative to the *in vitro* F_1_ and founding parents. (A) Field isolates, *in vitro* F_1_, *in vitro* selfs, and consensus parental genotypes plotted along the first two principal components (PCs). Each year is represented by a different color, with the A1 and A2 parental isolates indicated by blue and red, respectively. Shapes indicate the mating type of each isolate, with triangles (A1) and circles (A2). (B) A PCA of only the field isolates, with color and symbol scheme consistent with (A). (C) Pairwise Fst for comparisons between sample years and the *in vitro* F_1_ represented by a heat map, with more positive F_ST_ values increasingly red. A border indicates that the pairwise F_ST_ value was significantly different from 0, as tested by 1000 random SNP permutations.

To explore structure exclusively within the field population, we performed PCA on only the field isolates. Along PC1, isolates from 2012-13 were differentiated from prior year isolates (Figure 2B). Whereas, PC2 described differentiation within and between years.

To assess the variance in allele frequencies between years, we estimated pairwise F_ST_ (Weir and Cockerham, 1984) between years, where each year was defined as a distinct population. All pairwise comparisons were significantly greater than zero with the exception of 2009 versus 2010. Small F_ST_ estimates for comparisons between 2009, 2010, 2011 and the *in vitro* F_1_ indicated minimal variation in allele frequencies between these years. The greatest differences were observed between years 2012 and 2013 compared to 2009, 2010 and the *in vitro* F_1_ populations (Figure 2C), consistent with the PCA results. In addition, years 2012 and 2013 were also significantly differentiated from each other (F_ST_=0.027).

### Inbreeding in the field population

To quantify changes in inbreeding in the closed, field population, we estimated the individual inbreeding coefficient (F) for each isolate. While F does not directly measure identity-by-descent (IBD), it is highly correlated with IBD estimates in empirical and simulated data sets with relatively large numbers of markers (Kardos et al., 2015), particularly in highly subdivided, small populations (Balloux et al., 2004), such as the population under study. And, in a closed, biparental population, heterozygosity is directly proportional to the degree of inbreeding (Wright, 1921). Negative F estimates correspond to heterozygote excess relative to Hardy-Weinberg expectations for a reference population, defined here as the *in vitro* F_1_. Positive F values indicate heterozygote deficiency.

First, to establish expectations for a known F_1_ population, we assessed the F distribution in the *in vitro* F_1_. The *in vitro* F_1_, with a mean F of-0.366, was more heterozygous than the founding parents (average F across replicates=−0.007 (A1) and −0.183 (A2); Figure 3A). In contrast to the unimodal *in vitro* F_1_, the field population had a bimodal F distribution, with one peak approximately centered at the *in vitro* F_1_ mean, and a second peak centered at a less negative F value. This second peak indicated that inbreeding was occurring in the field population.

**Figure 3.**
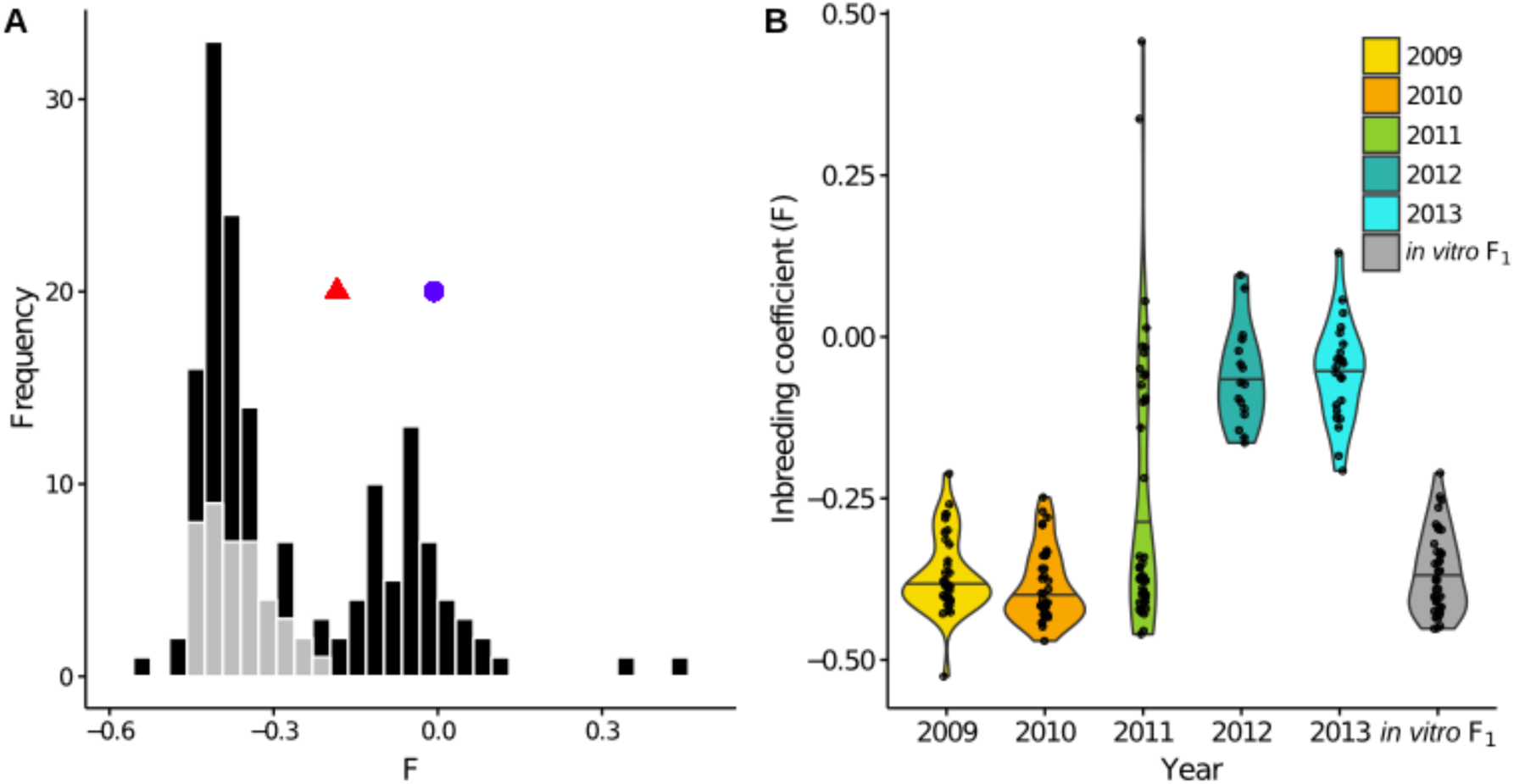
Generational shift in the field population. (A) Superimposed histograms of the individual inbreeding coefficient (F), estimated from 6,916 SNPs, in the *in vitro* F_1_ (gray) and field population (black). The *in vitro* F_1_ were more heterozygous than the founding parental isolates, corresponding to more negative F values, indicated by a blue circle (A1 parent) and red triangle (A2 parent). In contrast, the field population exhibited a bimodal F distribution. (B) Distributions of F by year represented by violin plots, with each year represented by a distinct color and individual data points overlaid. The long upper tail of the 2011 distribution is driven by two field selfs.

To dissect the bimodal shape of the field distribution, we analyzed F for each year separately. Both for 2009 and 2010, the distributions were unimodal with F means not significantly different from the *in vitro* F_1_ mean (pairwise t-test; *P*-values=1.0; Figure 3B). For years 2012 and 2013, distributions were also unimodal, but had F means significantly less negative than the *in vitro* F_1_ (*P*-values<0.0001). Year 2011 had a bimodal F distribution.

To interpret the effect of changes in inbreeding on genotypic and allele frequencies with time, we analyzed both SNP heterozygosity and MAF distributions for each year. In a biparental cross, clear expectations for these quantities in the F_1_ generation makes them informative in distinguishing F_1_ from inbred generations. Specifically, in the F_1_ generation, sites should segregate with a MAF of either 0.25 (for a cross of *Aa* x *AA)* or 0.5 (for *Aa* x *Aa* and *AA* x *aa*), and population heterozygosity should be 50% or 100% at each SNP. In the F_2_ generation, i.e. a population derived from a single generation of inbreeding, MAF should remain constant, whereas heterozygosity should decline. Our results showed that the *in vitro* F_1_, 2009, and 2010 behaved in accordance with expectations for a predicted F_1_; the heterozygosity distributions had peaks centered at approximately 50% and 100% (Figure 4A), and the MAF distributions had peaks at 0.25 and 0.5 (Figure 4B). In contrast, MAF and heterozygosity distributions in 2012 and 2013 were not consistent with F_1_ expectations, in that we no longer observed obvious peaks (Figure 4). While genotypic frequency shifts in 2012 and 2013 indicated presence of inbreeding and deviation from F_1_ expectations, changes in the MAF distribution also denoted that these were likely not canonical F_2_ populations. Discrete generations are implicit in a F_2_, therefore deviation from F_2_ expectations may be attributed to violation of this assumption.

**Figure 4.**
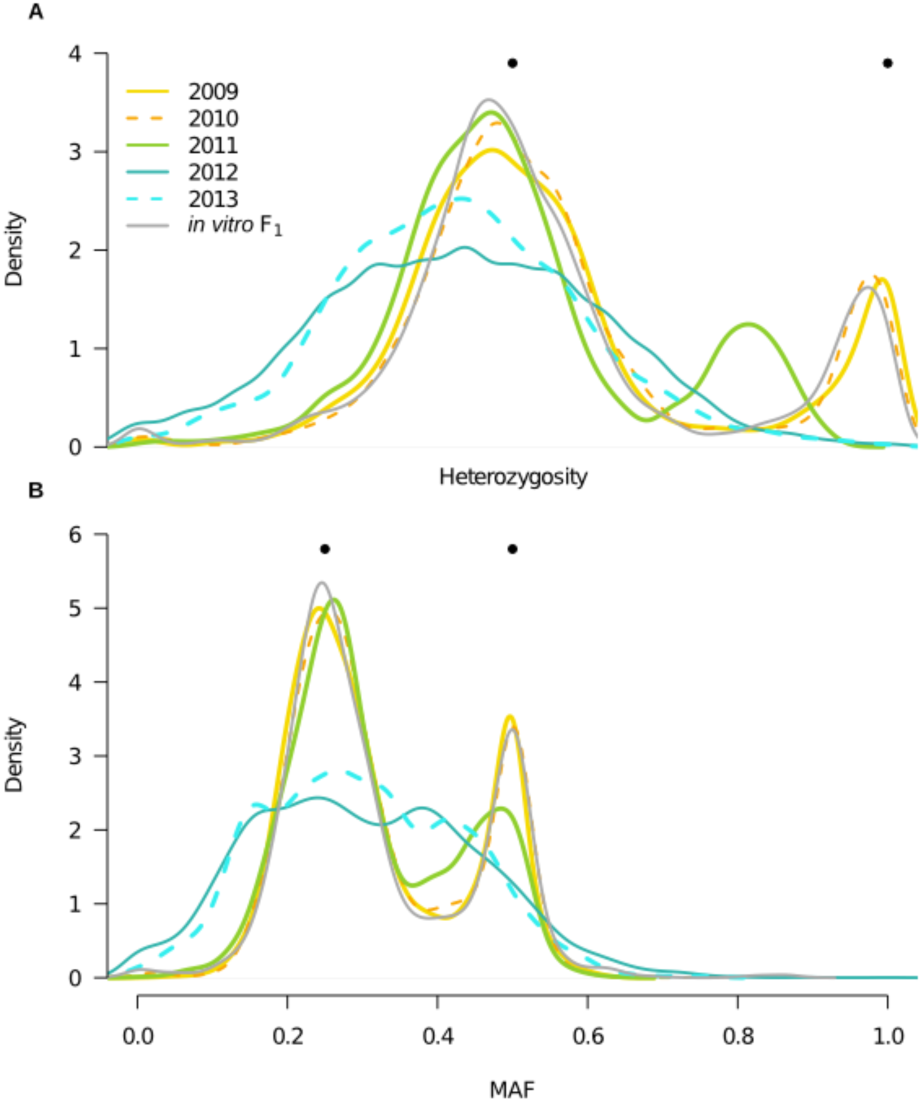
Year heterozygosity and allele frequency (MAF) distributions. Filled, black circles indicate expectations for population heterozygosity and MAF in a theoretical F_1_ population. (A) Distributions of the proportion of heterozygous individuals per SNP (*n*=6,916) for each year and the *in vitro* F_1_, represented by kernel density estimates, with color corresponding to year. Bimodal distributions in the *in vitro* F_1_ and years 2009-10 are consistent with expectations for the F_1_ generation, whereas unimodal distributions in 2012-13 indicate presence of inbreeding. A shift in the bimodal distribution of 2011, indicates the mixed outbred and inbred composition of this year. (B) MAF distributions, where the minor allele is defined based on the frequency in the total field population, for each year and the *in vitro* F_1_, with color designations the same as in (A).

Finally, in 2011, both heterozygosity and MAF distributions were bimodal, as in an F_1_, but with reduced heterozygosity and deviation in allele frequencies relative to the *in vitro* F_1_ and prior years (Figure 4). These shifts in 2011 suggested coexistence of both F_1_ and inbred isolates (i.e. non-F_1_ isolates) in this year, consistent with the bimodal 2011 F distribution (Figure 3B).

### Selfing in the laboratory and field

In addition to quantifying inbreeding (defined as inter-mating between related isolates), we also estimated the incidence of self-fertilization in the biparental, field population. The frequency at which *P. capsici* reproduces through self-fertilization in either field or lab conditions is unknown (Dunn et al., 2014). Given the limited prior evidence of selfing in *P. capsici,* we first confirmed that the three putative *in vitro* selfs were indeed the product of self-fertilization by the A1 parent, as hypothesized by Dunn et al. (2014). To this end, we distinguished the *in vitro* selfs from the *in vitro* F_1_ by four features: 1) Clustered with the A1 parent in PCA (dark blue circles in Figure 2A); 2) Alleles shared IBS disproportionately with the A1 versus A2 parent; 3) Heterozygosity approximately 50% of the A1 parent; and 4) significantly higher inbreeding coefficients relative to the F_1_ (>3 standard deviations (s.d.) from the mean; Table 1).

**Table 1.** Selfed isolates in the *in vitro* and biparental field populations in terms of heterozygosity, Mendelian errors (MEs), and alleles shared identity-by-state (IBS) with either founding parent.

|  | Consensus parental |  | *in vitro* selfs |  |  | Putative field selfs |  |
| --- | --- | --- | --- | --- | --- | --- | --- |
| Statistic | A1 | A2 | 68 14 | 68 19 | 68 27 | 11PF 21A | 11PF 26A |
| Individual heterozygosity<sup>1</sup> | 0.41 | 0.48 | 0.25 | 0.21 | 0.22 | 0.22 | 0.27 |
| $F^2$ | -0.01 | -0.18 | 0.38 | 0.48 | 0.46 | 0.46 | 0.34 |
| MEs<sup>3</sup> | 0.19 | 0.19 | 0.25 | 0.28 | 0.27 | 0.27 | 0.21 |
| IBS with the A1 parent<sup>4</sup> | 1.00 | 0.47 | 0.91 | 0.89 | 0.90 | 0.65 | 0.66 |
| IBS with the A2 parent<sup>4</sup> | 0.47 | 1.00 | 0.45 | 0.44 | 0.44 | 0.62 | 0.65 |

<sup>1</sup> Proportion of heterozygote, non-missing sites per individual
<sup>2</sup> Individual inbreeding coefficient (F)
<sup>3</sup> Proportion of Mendelian errors (MEs) per individual (corrected for outlier SNPs), relative to the consensus parental genotypes
<sup>4</sup> Identity-by-state (IBS) of each isolate with the A1 or A2 parental consensus genotype

Having shown that generalized expectations for selfing applied to *P. capsici,* we utilized extreme heterozygote deficiency as an indicator of selfing in the field. As, in the field context, the first three aforementioned selfing features were inapplicable because the progenitor of a selfed isolate in the field was not *a priori* known. We observed that two of the 2011 field isolates were F outliers (>3 s.d. from the mean) with respect to the inbred field contingent distribution (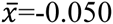, *s*=0.12). We classified these two A1 field isolates as field selfs (Table 1). Lack of disproportionate IBS of the field selfs with either founding parent denoted that these isolates were not the product of self-fertilization by either founding parent. Therefore, we observed selfing in the *in vitro* and field populations at frequencies of 3/46 (6.5%) and 2/159 (1.26%), respectively, denoting minimal incidence of selfing in both lab and field scenarios.

### Classifying F_1_ versus inbred isolates in the field using Mendelian errors (MEs)

Based on the above results, we hypothesized that 2009-10 were comprised of mainly F_1_, 2012-13 inbred, and 2011 a mixture of both F_1_ and inbred isolates. However, we had heretofore not verified that each year was homogeneous with respect to F_1_ and inbred composition. To quantify the number of F_1_ isolates, we used the fact that the genotypes of the founding parents were known to calculate an additional individual summary statistic, the proportion of Mendelian errors (MEs). A ME is defined as a genotype inconsistent with the individual being an F_1_ derived from specific parents (Purcell et al., 2007), here, the A1 and A2 founders. Commonly, MEs have been used to detect genotyping and experimental errors in SNP data sets where pedigree information is known (Purcell et al., 2007). The expectation is that a true F_1_ individual should have very few MEs, a postulate we applied to assess whether each field isolate belonged to the F_1_ generation.

Initial ME estimates revealed both randomly distributed and clustered ME-enriched SNPs. In S1 Text, we show that clustered ME-enriched SNPs corresponded to inferred mitotic LOH events in the parental isolates in culture. After removing all ME-enriched SNPs (*n*;=848), mean MEs per isolate for the *in vitro* F_1_ and field F_1_ subpopulations were 1.38% and 0.98%, below our estimated genotyping error rate of approximately 1.5%.

Akin to F, the proportion MEs per individual is a function of genotypic frequencies. Therefore, it was not surprising that year distributions of the ME statistic were consistent with F, with increased MEs in years 2012-13 (Figure 5A). Because asexual propagules do not survive the winter (Babadoost and Pavon, 2013; Hausbeck and Lamour, 2004), it can be assumed that all F_1_ isolates in the field, in any year, were derived from oospores in the year of the initial field inoculation (2008). Applying a threshold of 5.58% MEs (3 s.d. from the *in vitro* F_1_ mean) to characterize F_1_ versus non-F_1_, we observed exclusively F_1_ in 2009-10, a mixture of F_1_ and inbred isolates in 2011 (ratio of F_1_ to inbred=29:16) and all inbred isolates in 2012-13 (Figure 5B). As such, F_1_ dominated in 2009-11, demonstrating that oospores were viable and pathogenic for at least three years.

**Figure 5.**
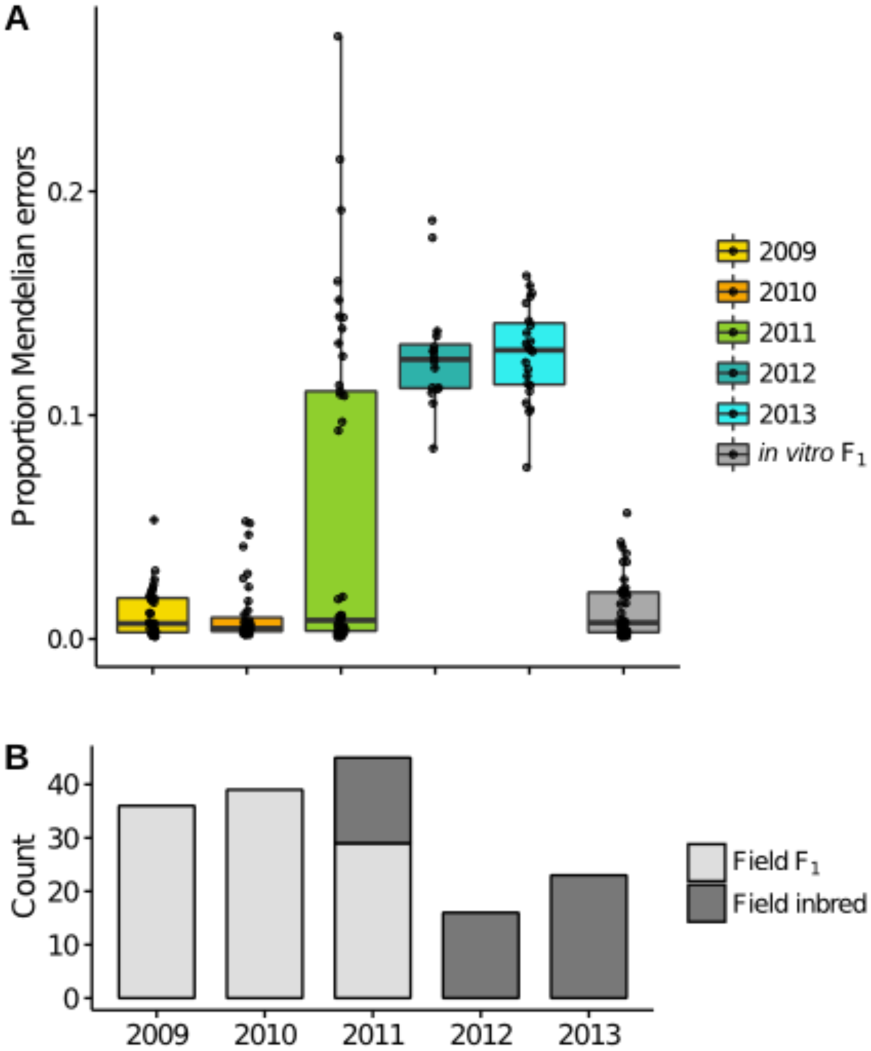
Mendelian errors (MEs) distinguish F_1_ and inbred isolates in the field. (A) Boxplots of the proportion of MEs per individual for each year are consistent with the inbreeding coefficient trend, with a bimodal distribution in 2011, and increased MEs in later years. (B) classification of each isolate based on the proportion of MEs, with counts of field F_1_ (light gray) and field inbred (dark gray) for each sample year.

When the inbred isolates were removed from the 2011 data, the MAF distribution for 2011 was consistent with F_1_ expectations (Supplementary Figure S5). Concurrent observation of both F_1_ and inbred isolates in a single year (2011) provided direct evidence of overlapping generations in the field population, supporting overlapping generations as contributing to deviation from F_2_ expectations in the inbred 2012 and 2013 years.

In addition, the ME estimates allowed us to pool isolates from separate years to define sub-populations, the field F_1_ (*n*=104) and the field inbred (*n*=53; excluding the field selfs), for subsequent analyses. As in the total field population, A2 isolates were overrepresented in both the field F_1_ and inbred subpopulations (A1:A2=43:61 and 21:32, respectively).

### Regions of differentiation between generations in the field population

The generational transition in the field population from F_1_ to inbred was accompanied by changes in the MAF distribution (Supplementary Figure S5), implying the biased transmission of alleles to generations beyond the F_1_. To identify which SNPs drove this allele frequency shift, we performed a genome-wide Fisher’s Exact test of allele frequency differences between the field F_1_ and field inbred subpopulations. We collectively analyzed these two subpopulations, rather than compare allele frequencies between years, due to the presence of overlapping generations, which complicate interpretation of temporal dynamics (Jorde and Ryman, 1995). From this analysis, we observed several regions of differentiation between these subpopulations (Figure 6; see Supplementary Table S5 for coordinates).

**Figure 6.**
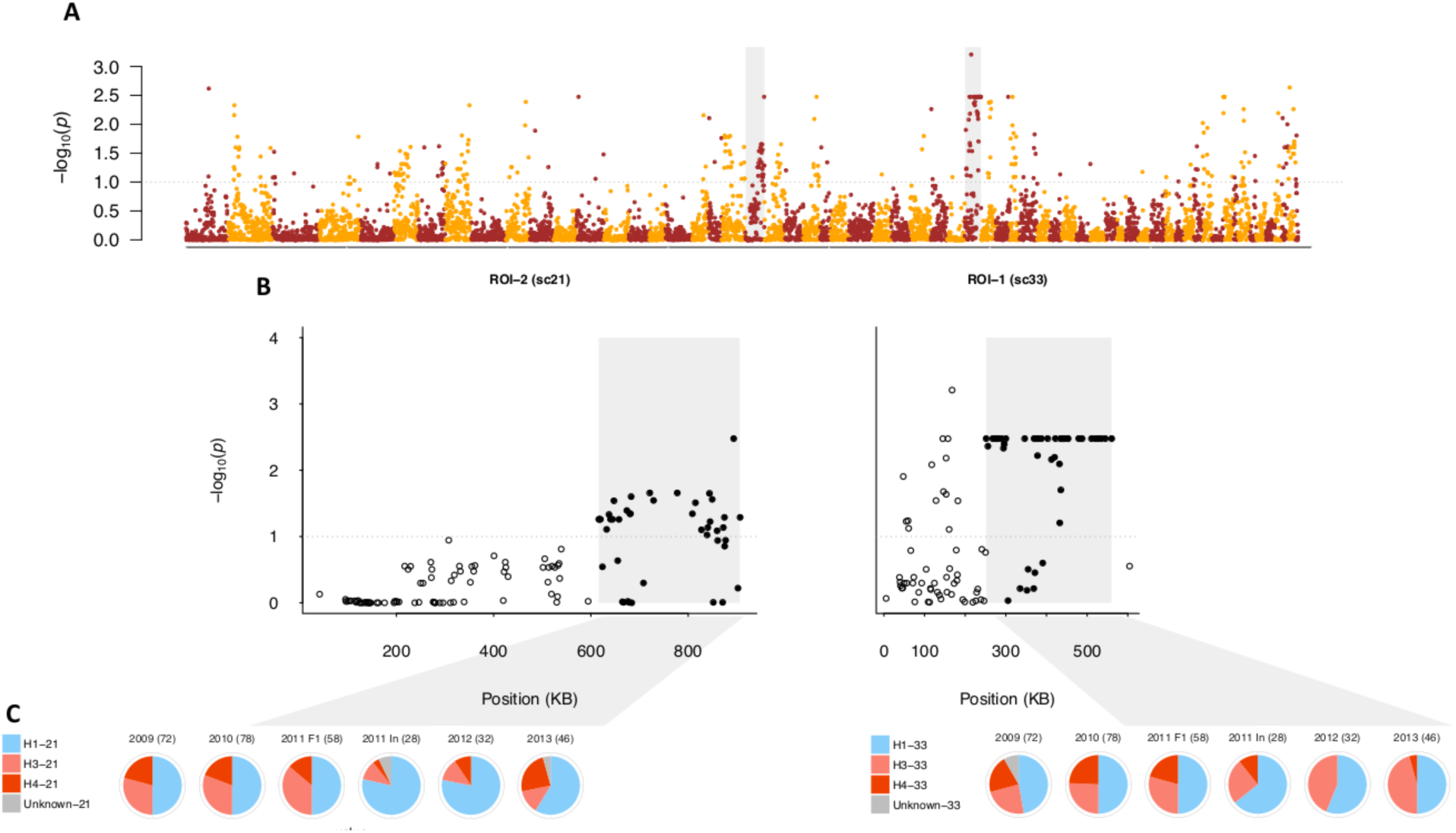
Regions of differentiation between the field F_1_ and inbred subpopulations. (A) Negative log_10_-transformed, false-discovery rate (FDR) adjusted *P*-values from the genome-wide test of allele frequency differences between the field F_1_ and inbred subpopulations, ordered by physical position. The gray dotted lines in (A) and (B) indicate the significance threshold (α=0.10). Color alternates by scaffold. The shaded gray boxes indicate the SNPs in scaffolds 21 and 33 corresponding to ROIs 2 and 1, respectively. (B) Same as (A) except that *P*-values are shown only for scaffolds 21 and 33. Here, gray boxes denote the sub-region within each scaffold defined as a ROI. Closed, black circles indicate SNPs within each ROI, whereas open, black circles indicate SNPs outside of the ROI. (C) Pie charts represent the haplotype frequencies found in each year (with 2011 separated into F_1_ and inbred (In) isolates), with the number of sampled chromosomes noted for each year. Blue corresponds to the single A1 founding parental haplotype, shades of red to the two A2 founding haplotypes, and gray to undesignated haplotypes in each ROI (see Methods)

First, we focused on the region with the most highly differentiated SNP, referred to as region of interest 1 (ROI-1; Figure 6B). Of the 94 SNPs spanned by ROI-1, 44% were among SNPs in the top 2% of loadings for PC1 in the field PCA, showing that this region was correlated with differentiation in the field population. To assess the relationship between allele frequency changes and parental haplotype frequencies, we locally phased all isolates using a deterministic approach (see Methods). Haplotypes in ROI-1 (H1a, H3a, and H4a) were defined based on the sub-region (251,367-560,094 bp) which contained the majority of significantly differentiated SNPs (44 out of 52 SNPs) and formed a LD block (Supplementary Figures S11 and S14).

Segregation among the F_1_ isolates in each year (2009 to 2011) followed the F_1_ expectation of a 2:1:1 ratio of H1:H3:H4 haplotypes (*x*^2^ test; *P*-values=0.91, 0.99, 0.65, respectively). In contrast, in 2011 (inbred isolates only), 2012, and 2013, we observed lower frequencies of H4 and higher frequencies of H3 relative to the field F_1_ subpopulation (Figure 6C). The decline in H4 frequency from 22.12% in the field F_1_ to 4.72% (and corresponding increases in H3 and H1) in the field inbred drove allele frequency changes in ROI-1 (Supplementary Table S6). Because the H4 sequence was most distinct from the other haplotypes, the reduction in H4 frequency, along with inbreeding, resulted in declines in heterozygosity in ROI-1. Consistent reductions in H4 frequency among inbred isolates in 2011-13 compared to F_1_ isolates in prior years, provided strong evidence for the influence of selection. However, absence of H4 in year 2012 is very likely an artifact of smaller sample size in this year.

We next focused on a region in scaffold 21, defined as ROI-2, with the highest density of significantly differentiated SNPs (67%; Figure 6B). In ROI-2, as in ROI-1, only three haplotypes segregated in the field population (Supplementary Table S7). While not significant (at a=0.05), segregation among the F_1_ isolates in each year (2009 to 2011) deviated from the F_1_ expectation of a 2:1:1 ratio of H1:H3:H4 haplotypes (*χ*^2^ test; *P*-values=0.61, 0.35, and 0.05, respectively), primarily attributed to higher H3 versus H4 haplotype frequency in the field F_1_ (*χ*^2^ test; *P*-value<0.01). A decline in frequency of the A2 parent haplotype, H3, by 19.47% and an increase in the Al parent haplotype, H1, by 19.81% drove allele frequency changes (Figure 6C and Supplementary Table S7). While the frequency of H3 and H4 oscillated among inbred isolates in 2011-13, the H1 haplotype frequency was consistently higher than in the field F_1_. In addition, we observed a high frequency of homozygous H1 genotypes (53%), whereas the H3 and H4 haplotypes were not observed in the homozygous state in the field inbred subpopulation, contrary to expectations (Supplementary Table S7).

To posteriorly assess the significance of changes in allele frequency in ROIs 1 and 2, we compared the median F_ST_ value for significantly differentiated SNPs in each of these regions to the genome-wide SNP F_ST_ distribution, where F_ST_ was defined as in (Lewontin and Krakauer, 1973). Assuming that drift acts equally throughout the genome, extreme deviations in F_ST_ provide evidence for selection (Lewontin and Krakauer, 1973). Median observed changes in allele frequency in ROIs 1 and 2 were in the in the 97^th^ and 98^th^ percentiles, respectively, relative to genome-wide F_ST_, showing that allele frequency changes in these regions vastly exceeded the genome-wide average.

### Heterozygosity declines are slower in the mating type region

To investigate whether the mating system was a direct driver of differentiation in the field population, we first identified mating type associated SNPs using a Fisher’s exact test of allele frequency differences between isolates of opposite mating types in the field F_1_ (*n*_A1_=43 and *n*_A2_=61; Supplementary Figure S15). Most of the 184 significantly differentiated SNPs were in sub-regions of scaffolds 4 (37%) and 27 (43%), with additional differentiated SNPs in sub-regions of scaffolds 2, 34, and 40 (Figure 7 and Supplementary Table S8). All scaffolds containing significantly associated SNPs were in linkage group 10, consistent with a prior study (Lamour et al., 2012), and supporting presence of a single mating type determining region in *P. capsici,* as posited for *P. infestans* and *P. parasitica* (Fabritius and Judelson, 1997). SNPs in these five sub-regions comprised 20.29% of SNPs with elevated PC loadings (top 2%) in the PCA of only the field isolates, compared to 5.10% genome-wide, denoting that these SNPs were disproportionately correlated with differentiation in the field population.

**Figure 7.**
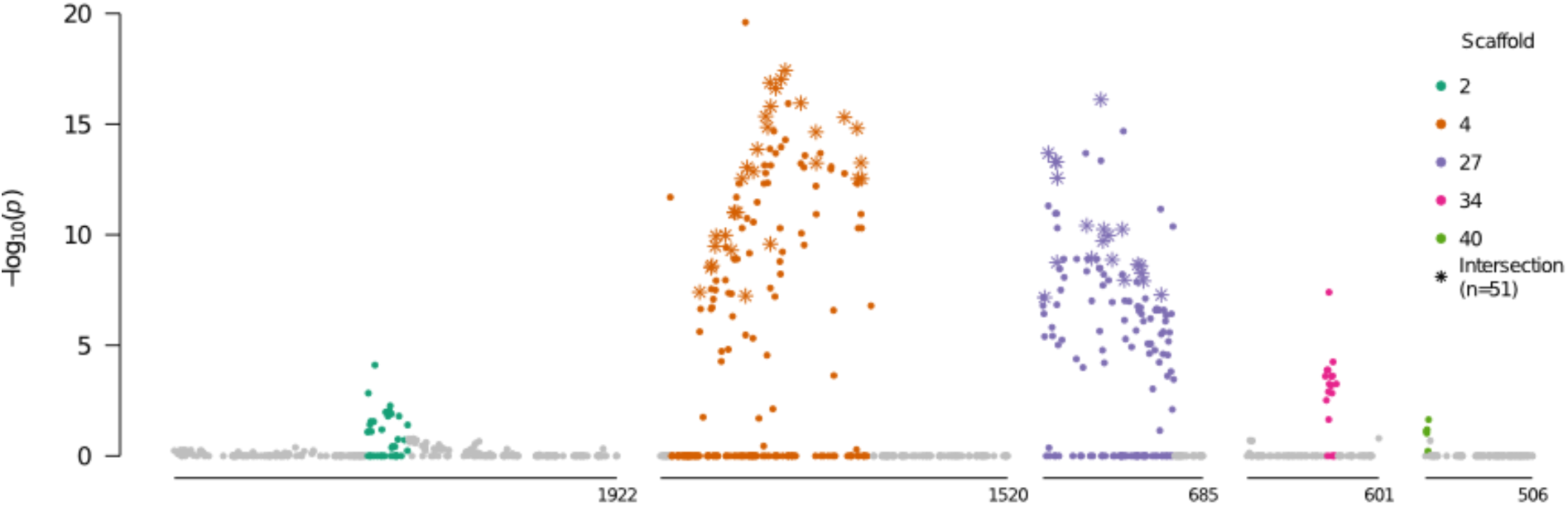
Allele frequency differences between isolates of opposite mating types. Negative log_10_-transformed *P*-values, adjusted for multiple testing, from the fisher’s exact test of allele frequency differences between A1 and A2 isolates in the field F_1_, plotted against physical position, for scaffolds with significantly differentiated regions (see Supplementary Table S8 for coordinates). Colored SNPs were within the bounds of the minimum and maximum significant SNPs in each scaffold containing at least two significantly associated SNPs within 200 kb. Stars indicate the SNPs which were significant in tests of allele frequency differences between mating types in both the field F_1_ and inbred subpopulations (Supplementary Text). All SNPs above the gray horizontal line were significant after the FDR correction (α=0.1).

At 98.30% of the *AA* x *Aa* SNPs associated with mating type in the field F_1_, the A2 parent was heterozygous (*Aa*) and the A1 parent was homozygous (*AA*). As such, heterozygosity in the field progeny at these SNPs was attributed to inheritance of the minor allele (*a*), descendent originally from the A2 parent. Therefore, segregation of the A2 but not A1 parental haplotypes was predominantly associated with mating type in the field F_1_.

We defined the mating type region (MTR) as consisting of genomic tracts encompassed by the minimum and maximum significant SNPs in scaffolds 4 and 27, which comprised 1.42 of the 1.64 Mb spanned by the five sub-regions, and contained 81% of the significantly differentiated SNPs. While we refer to a singular MTR, this was not intended to imply physical linkage between these two scaffolds. Based on the 293 SNPs in the MTR, the PCA of all isolates (*in vitro* and field; *n*=203) showed incomplete differentiation according to mating type (Supplementary Figure S16).

To assess changes in heterozygosity in the MTR, we compared the heterozygosity distributions of the field F_1_ (*n*_A1_=43 and *n*_A2_=61) and inbred (*n*_A1_=21 and *n*_A2_=32) isolates in the MTR to the respective genome-wide distributions (see Methods). Observed heterozygosity in the field F_1_ in the MTR was not centered at a significantly greater mean than the field F_1_ genome-wide distribution (one-sided Wilcoxon rank-sum test, all *P*-values>0.87; Supplementary Figure S17). In contrast, observed heterozygosity in the field inbreds in the MTR was shifted towards a greater mean relative to the field inbred genome-wide distribution (all *P*-values<0.005; Supplementary Figure S17). Therefore, in the field inbred subpopulation, heterozygosity declines were less appreciable in the MTR compared to the rest of the genome. In addition, we found that heterozygosity in the MTR was significantly higher than the rest of the genome for both the A1 and A2 isolates in the field inbred subpopulation (all *P*-values<10^−4^ and 0.003, respectively; Supplementary Figure S17), but not in the field F_1_ (all *P*-values>0.99 and 0.65, respectively; Supplementary Figure S17). Yet, heterozygosity in the MTR did not significantly exceed HWE expectations for A1 inbred isolates, as observed in the A2 inbred isolates, and both mating types in the field F_1_ (Supplementary Figure S18). These results were replicated when the A2 inbred isolates were down-sampled to the A1 inbred sample size (data not shown).

To further dissect the genetic dynamics of mating type in the field population, we tracked the allele frequencies of markers that were heterozygous in one parent and homozygous in the other parent (*AA* x *Aa*). These markers are particularly informative because the origin of the *a* allele can be unambiguously assigned to the heterozygous parent. Specifically, we calculated the frequency of the parental tagging allele (*p*_*a*_) in the parents, the field F_1_, and the field inbreds at each of the *AA* x *Aa* SNPs in the MTR (*n*_*AA*x*Aa*_=206), for each mating type separately.

For A2 tagging SNPs (*n*=98), i.e. SNPs heterozygous in the A2 founding parent, we observed two classes of markers: those with *p_a_* of approximately 0.5 in the A2 and 0.0 in the A1 field F_1_ isolates, and the opposite scenario (Figure 8A-B; see Methods). In the First case (*n*=49), differences in *p_a_* between mating types were maintained in the field inbred subpopulation (Figure 8A). Whereas, in the second case (n=49), the difference in *p*_*a*_ between mating types narrowed (Figure 8B). In contrast to the A2 parent tagging SNPs, markers heterozygous in the A1 parent and homozygous in the A2 parent (*n*=108) predominantly followed a single pattern (Figure 8C). These markers were at approximately equal allele frequencies in both mating types in the field F_1_, but slightly diverged in frequency in the field inbred subpopulation. These three distinct segregation patterns were consistent with the association of presence/absence (P/A) of one of the A2 founding haplotypes in association with mating type.

**Figure 8.**
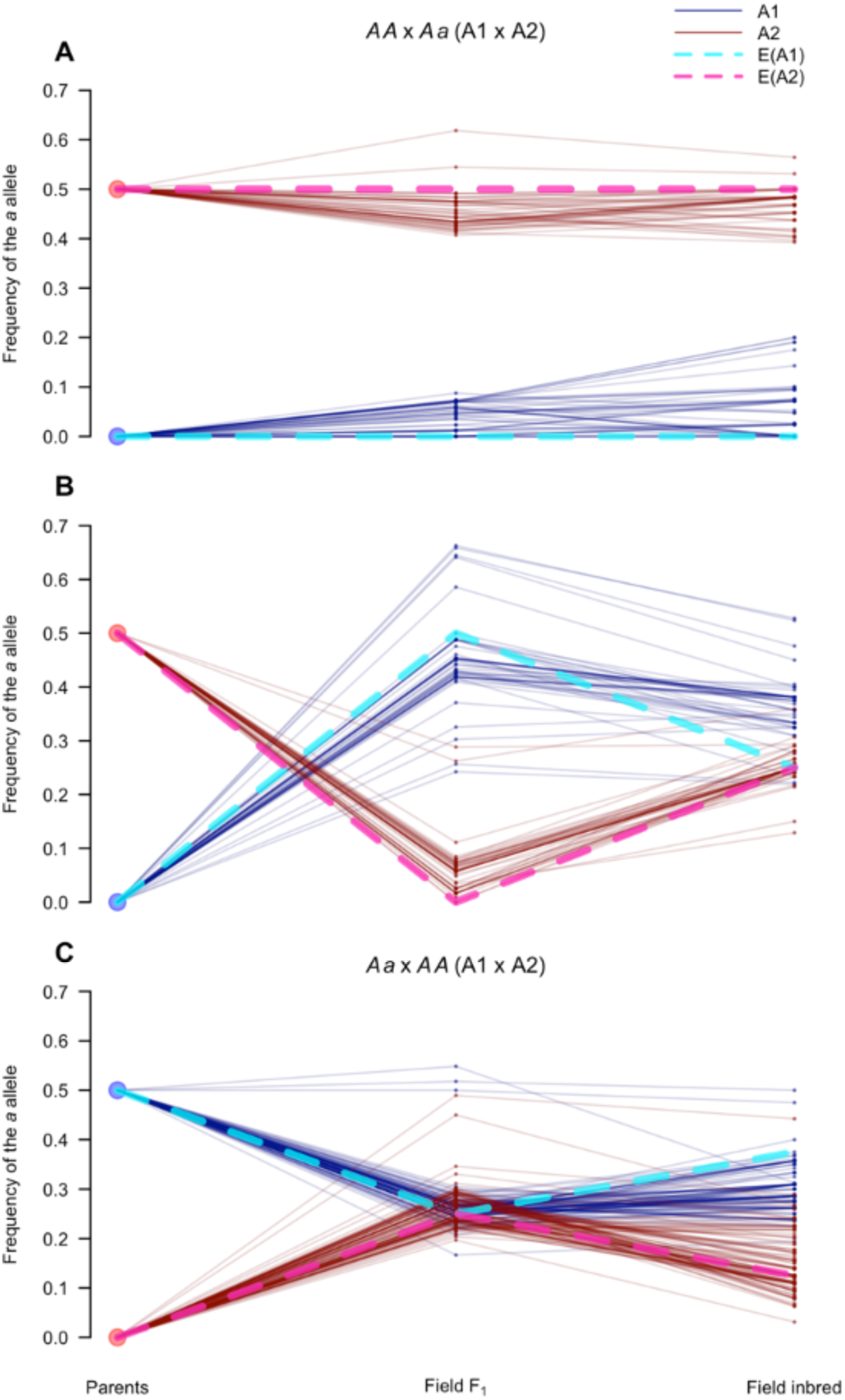
Segregation of SNPs in the mating type region follow expectations for sex-linked loci. Frequency of the *a* allele (*p*_*a*_), for *AA* x *Aa* and *Aa* x *AA* (A1 x A2) markers in the mating type associated sub-regions of scaffolds 4 and 27, defined as the mating type region (MTR). Each parallel coordinate plot (A-C) tracks *p*_a_ at three time points (parents, field F_1_, and field inbred) in the A1 (blue solid lines) and A2 (red solid lines) isolates for: (A) *AA* x *Aa* markers (*n*=49) with *p*_a_>0.3 in the A2 and *p*_a_<0.3 in the A1 field F_1_ isolates; (B) Remaining *AA* x *Aa* markers (*n*=49); and (C) All *Aa* x *AA* markers. Expectations for sex-linked loci, indicated by dotted lines, assuming the A1 and A2 mating types behave like the homogametic (light blue) and heterogametic (pink) sexes, respectively, when: A) the *a* allele is in the A2 determining haplotype (i.e. Y analog); B) the *a* allele is in the non-A2 determining haplotype (i.e. X analog); and C) the *a* allele is in either of the non-A2 determining haplotypes (i.e. X analog).

While the structural basis of mating type determination in *P. capsici* is not known, observed segregation patterns in the mating type region resemble those of an XY system, where P/A of the Y determines sex. Therefore, as frame of reference, we derived expectatinos for sex-linked loci in an XY system, i.e. loci conserved between both sex chromosomes (Allendorf et al., 1994; Clark, 1988), assuming that the A2 parent corresponded to the heterogametic sex. Using this model, expectations (blue and pink dotted lines in Figure 8) closely matched the observed *p*_*a*_ trajectories in all three cases (A-C), further supporting the association of one of the A2 haplotypes with mating type determination.

## Discussion

To study the temporal genetic dynamics of *P. capsici* in response to a severe bottleneck, we SNP genotyped at high-density 232 isolates collected in years 2009-13 from a closed, biparental field population founded in 2008, in Geneva, NY (Dunn et al., 2014). This experimental population parallels the infection scenario of a natural *P. capsici* epidemic, where a limited number of pathogen strains are thought to found a subsequently isolated population (Dunn et al., 2014; Lamour et al., 2012). Using filtered GBS data, we identified 159 unique field isolates and obtained 6,916 high quality SNPs with high sequencing depth (∼20X coverage), low missing data, and over 97% reproducibility of genotype calls, distributed throughout the genome. With these data, we assessed temporal heterozygosity and allele frequency changes in the biparental population, representing the only controlled, multi-year genomic field study of a plant pathogen to date.

With knowledge of the parental genotypes and assuming simple Mendelian inheritance, we developed a threshold to detect F_1_ field isolates based on the incidence of MEs in the *in vitro* F_1_ progeny. Our results showed that both field and *in vitro* F_1_ progeny were characterized by individual heterozygosity in large excess of Hardy-Weinberg expectations, explained by the fact these isolates were descendent from only two parents. With small numbers of parents, the probability of allele frequency differences between opposite sexes (here, mating types) increases, consequently resulting in deviation from HWE among the progeny (Balloux, 2004; Luikart and Cornuet, 1999; Pudovkin et al., 1996; Robertson, 1965).

Over time, the field population underwent a generational shift, transitioning from F_1_ in 2009-10, to mixed generational in 2011, and ultimately all inbred in 2012-13. Presence of exclusively F_1_ in 2009 suggests that the vast majority of oospores formed in the founding year (2008) were F_1_. As oospores require a dormancy period of approximately one month (Dunn et al., 2014; Satour and Butler, 1968), it is not surprising that there was insufficient time to produce multiple generations in the founding year. The presence of only F_1_ and no inbred isolates in 2010, however, cannot be similarly explained. Rather, abundant sexual reproduction in the founding year, coupled with a lower rate in 2009, may have led to disproportionate presence of F_1_ oospores (from 2008) surviving in the soil and germinating in 2010. Year 2011, where both inbred and F_1_ isolates were observed in the field, signified a generational shift in the population. The absence of F_1_ in the following years (2012-13) is consistent with previous reports of oospore declines in viability over time (Bowers, 1990), and negligible oospore survival after four years in field conditions (Babadoost and Pavon, 2013). While we did not quantify disease incidence in the field, observation of predominantly F_1_ isolates in 2010-11 suggests that highly productive years contributed disproportionately to population structure, in accordance with theoretical predictions for populations in which sexual propagules require a dormancy period, e.g. plant species with seed banks (Nunney, 2002; Templeton and Levin, 1979). As a consequence, heterozygosity did not immediately decline in the second year of the field population, similarly consistent with the delayed attainment of equilibrium genotypic frequencies attributed to seed bank dynamics (Templeton and Levin, 1979).

Approximate equilibrium genotypic frequencies were not observed in the field population until the fourth year (2012). Here, a single large increase in homozygosity in the total population was consistent with cycles of inbreeding beyond a theoretical F_2_ resulting in less appreciable declines in heterozygosity relative to the prior generation (Wright, 1921). However, two excessively homozygous field isolates, identified as field selfs, significantly deviated from this trend. Given this low frequency of selfing, we conclude that *P. capsici* behaved essentially as an obligate outcrossing species in the biparental field population. Occurrence of selfing in *P. capsici* is consistent with a previous report of oospore induction when strains of opposite mating types were separated by a membrane (Uchida and Aragaki, 1980), but contradicts previous studies which found no evidence for self-fertilization under *in vitro* conditions (Hurtado-Gonzales and Lamour, 2009; Lamour et al., 2012). As a single generation of self-fertilization reduces heterozygosity by approximately 50% in the progeny, minimal incidence of selfing delayed heterozygosity declines in the field population, as described for hermaphroditic plant species (Balloux, 2004).

In addition, while we observed three A1 parental selfs among the 46 *in vitro* progeny, we did not observe selfs derived from either founding parent in the field, despite the larger field F_1_ sample size. Field isolates were inherently selected for both viability and pathogenicity, as well as resilience to environmental factors, whereas, *in vitro* isolates were selected solely on viability in culture. Therefore, this result may reflect a fitness cost to self-fertilization, as observed in essentially all outcrossing species (Charlesworth and Charlesworth, 1987; Falconer and Mackay, 1996), manifest to a greater extent in the field versus laboratory conditions.

Given the potential fitness cost to self-fertilization in *P. capsici,* an increase in inbreeding may explain the allele frequency changes which accompanied the transition from an F_1_ to inbred population, as inbreeding presents recessive deleterious alleles in the homozygous state, rendering them subject to selection (Charlesworth, 2003; Kirkpatrick and Jarne, 2000). Simultaneously, inbreeding indirectly influences allele frequencies by decreasing the effective population size (N_e_) relative to the census population size, consequently amplifying the effects of genetic drift (Charlesworth, 2009). In addition to inbreeding, many other factors likely decreased N_e_, thereby increasing the influence of genetic drift: imbalanced sex (here, mating type) ratios (Charlesworth, 2009); clonal reproduction (Balloux et al., 2003); variation in reproductive success (Hartl and Clark, 2007); small population sizes (Hartl and Clark, 2007) suggested by lower genotypic diversity in 2012-13; and overlapping generations (Felsenstein, 1971). Conversely, minimal differentiation between 2009-11, denotes that oospore survival, in behaving like a seed bank, mitigates the aforementioned reductions in N_e_ by maintaining a reservoir of genetic variation in the soil (Hairston and De Stasio, 1988; Nunney, 2002; Templeton and Levin, 1979; Waples, 2006).

In contrast to the general trends described above, we characterized two regions (ROI-1 and ROI-2) that significantly deviated from the genome-wide distribution of allele frequency differences (median allele frequency change in the 97 percentile or greater) between the field F_1_ and inbred subpopulations. In these two regions, which presented only three segregating haplotypes, in contrast to the expected four, for heterozygous parents, we associated allele frequency changes with haplotype frequency shifts. Genetic drift may still explain these results, as drift has a larger effect in regions of low variation, i.e. with higher effective inbreeding coefficients corresponding to lower local N_e_ (Charlesworth, 2009; 2003). However, extreme changes in allele frequency are also suggestive of natural selection (Galtier et al., 2000; Lewontin and Krakauer, 1973). Here, observation of corresponding haplotype frequency shifts is consistent with hitchhiking or background selection having a large effect in inbred populations (Charlesworth, 2003), particularly with only a few generations. Alternatively, mitotic LOH, a phenomenon reported in the present study (see Supplementary Text) and in numerous *Phytophthora* species (Chamnanpunt et al., 2001; Grünwald et al., 2012; Kasuga et al., 2016; Lamour et al., 2012), may explain the observation of a disproportionate number of homozygous genotypes among inbred isolates in ROI-2. Evidence for mitotic LOH in numerous species, e.g. *Saccharomyces cerevisiae* (Magwene et al., 2011), *Candida albicans* (Forche et al., 2011), and the chytrid *Batrachochytrium dendrobatidis* (Rosenblum et al., 2013), supports the theoretical expectation that this process facilitates adaptation by interacting with selection to alter allele frequencies (Mandegar and Otto, 2007). Given the limited number of generations, we cannot unequivocally attribute these dramatic haplotype frequency shifts to selection. Furthermore, additional work is required to assess the role of these regions in pathogenicity and local adaptation.

While we observed a genome-wide increase in homozygosity in the field population due to inbreeding, reductions in heterozygosity in the identified mating type associated region were smaller relative to the genome for both A1 and A2 isolates. We show that this result is explained by persistent allele frequency differences between isolates of opposite mating types in the MTR. Maintenance of elevated heterozygosity in sex-linked regions has been attributed to differences in founding allele frequencies between sexes in several systems (Allendorf et al., 1994; Marshall et al., 2004; Waples, 2014). Further, *AA* x *Aa* SNPs associated with mating type in the field F_1_ were predominantly heterozygous in the A2 parent, implying that one of the A2 founding haplotypes was associated with mating type determination. Consistent with this result, segregation patterns for SNPs in the MTR resembled the behavior of loci in the pseudoautosomal (conserved) regions of heteromorphic sex chromosomes (e.g. XY or ZW; (Clark, 1988)), where the A2 parent corresponded to the heterogametic, male sex. These results suggest that in populations of *P. capsici* with few founders, heterozygosity in the MTR will be maintained despite inbreeding, proportional to LD between the mating type factor(s) and the rest of the genome.

These findings, which represent the first detailed genomic analysis of mating type in a *Phytophthora* species, are consistent with the existing models of heterozygosity versus homozygosity at a single locus as determinant of mating type (Fabritius and Judelson, 1997; Sansome, 1980). However, our analysis does not demonstrate that heterozygosity *per se* confers the A2 mating type, nor does our analysis preclude the presence of heteromorphic mating type chromosomes in *P. capsici.* We applied stringent SNP filters to obtain a high quality set of markers, likely discarding SNPs located in regions of structural variation (i.e. duplications, deletions, repeats). Indeed, early cytological work supports heterozygosity for a reciprocal translocation in association with mating type in *P. capsici* and numerous *Phytophthora* species, posited as a mechanism to suppress local recombination (Sansome, 1976). Given that chromosomal heteromorphism has arisen in diverse taxa as a consequence of suppressed recombination between sex-determining chromosomes (Bachtrog, 2013; Charlesworth, 2013), future studies will investigate the association of structural variation and recombination suppression with mating type determination in *P. capsici.*

## Materials and Methods

### Isolate and DNA collection

In 2008, a restricted access research field at Cornell University’s New York Agricultural Experiment Station in Geneva NY, with no prior history of Phytophthora blight, was inoculated once with two NY isolates of *P. capsici,* 0664-1 (A1) and 06180-4 (A2), of opposite mating types, as described in Dunn et al (2014). From 2009-13, the field was planted with susceptible crop species, and each year the pathogen was isolated from infected plant material, and cultured on PARPH medium (Dunn et al., 2014). Once in pure culture, a single zoospore isolate was obtained (Dunn et al., 2014), and species identity was confirmed with PCR using species specific primers, as previously described (Dunn et al., 2010; *Zhang* et al., 2008).

Isolates collected in 2009-12 were obtained from storage; isolates from 2013 were unique to this study and were collected from infected pumpkin plants (variety Howden Biggie). Single oospore progeny (*n*=46) from an *in vitro* cross between the founding parents were obtained from storage (Dunn et al., 2014). To revive isolates from storage, several plugs from each storage tube were plated on PARPH media. After less than one week, actively growing cultures were transferred to new PARP or PARPH medium.

Mycelia were harvested for DNA extraction as previously described (Dunn et al., 2010), except that sterile 10% clarified V8 (CV8) broth (Skidmore et al., 1984) was used instead of sterile potato dextrose broth. For each isolate, mycelia were grown in Petri plates containing CV8 broth for less than 1 week, vacuum filtered, and 90-110 mg of tissue were placed in 2ml centrifuge tubes and stored at −80°C until DNA extraction. DNA was extracted using the DNeasy Plant Mini kit (Qiagen, Valencia, CA) according to manufacturer’s instructions, except that mycelial tissue was ground using sterile ball bearings and a TissueLyser (Qiagen, Valencia, CA) as previously described (Dunn et al., 2010).

Mating type was determined as previously described (Dunn et al., 2010). Briefly, each isolate was grown on separate unclarified V8 agar with known A1 and A2 isolates, respectively. After at least one week of growth, the plates were assessed microscopically for the presence of oospores. For each trial, the A1 and A2 tester isolates were grown in isolation and on the same plate as negative and positive controls, respectively. We obtained mating type designations for isolates from years 2009-12 and the *in vitro* F_1_ from (Dunn et al., 2014).

### Genotyping

All DNA samples were submitted to the Institute of Genomic Diversity at Cornell University for 96-plex GBS (Elshire et al., 2011). In brief, each sample was digested with *Ape*KI, followed by adapter ligation, and samples were pooled prior to 100bp single-end sequencing with Illumina HiSeq 2000/2500 (Elshire et al., 2011). Sequence data for samples analyzed in this study are available at XXX (sftp site).

To validate experimental procedures, DNA samples from the parental isolates were included with each sequencing plate (except in one instance). The parental isolates were sequenced initially at a higher sequencing depth (12-plex). Genotypes were called for all isolates simultaneously using the TASSEL 3.0.173 pipeline (Glaubitz et al., 2014). This process involves aligning unique reads, trimmed to 64bp, to the reference genome (Lamour et al., 2012) and mitochondrial (courtesy of Martin, F., USDA-ARS) assemblies, and associating sequence reads with the corresponding individual by barcode identification to call SNPs (Glaubitz et al., 2014).

The Burrows-Wheeler alignment (v.0.7.8) algorithm bwa-aln with default parameters (Li and Durbin, 2009) was used to align sequence tags to the reference genome (Lamour et al., 2012). To reduce downstream SNP artifacts due to poor sequencing alignment, reads with a mapping quality <30 were removed. Default parameters were otherwise used in TASSEL, with two exceptions: 1) Only sequence tags present >10 times were used to call SNPs; and 2) SNPs were output in variant call format (VCF), with up to 4 alleles retained per locus, using the tbt2vcfplugin. Genotypes were assigned and genotype likelihoods were calculated as described in (Hyma et al., 2015).

### Individual and SNP Quality Control

Individuals with more than 40% missing data were removed from analysis. To mitigate heterozygote undercalling due to low sequence coverage, genotypes with read depth <5 were set to missing using a custom python script (available upon request). Subsequently, we utilized VCFtools version 1.14 (Danecek et al., 2011) to retain SNPs which met the following criteria: 1) Genomic; 2) <20% missing data; 3) Mean read depth ≥10; 4) Mean read depth <50; 5) Bi-allelic; and 6) Minor allele frequency (MAF)≥0.05. Additionally, indels were removed.

To remove isolates with likely ploidy variation, we assessed allele depth ratios for each isolate, where the allele depth ratio was defined as the ratio of the major allele to the total allele depth at a heterozygous locus (Li et al., 2015; Rosenblum et al., 2013; Yoshida et al., 2013). Allele depths were extracted from the VCF file, using a custom python script (available upon request) to analyze the distribution of allele depth ratios for each individual across all SNPs.

Post clone-correction (see below) and outlier removal, SNPs were further filtered as follows. SNPs with heterozygosity rates >90% among all isolates (clone-corrected and parental replicates) were removed and/or average allele depth ratios <0.2 or >0.8. Only SNPs within scaffolds containing more than 300 kb of sequence, covering ∼48 Mb (∼75% of the sequenced genome), were retained. We defined the minor allele as the least frequent allele in the clone-corrected field population.

Multiple sequencing runs of the parental isolates were used to define consensus genotypes for each parent using the majority rule. Sites where ≥50% of calls were missing or where disparate genotype calls were equally frequent were set to missing.

All filtering and analyses, if not otherwise specified, were performed in R version 3.2.3 (R Core Team, 2015) using custom scripts (available upon request).

### Identifying a clone-correction threshold

To establish a maximum similarity threshold to define unique genotypes, the genetic similarity of all sequencing runs of the parental isolates were compared. Similarity was defined as identity-by-state (IBS), the proportion of alleles shared between two isolates at non-missing SNPs. Parental replicates represented both biological (different mycelial harvests and/or independent cultures) and technical replicates (same DNA sample), thereby capturing variation associated with culture transfers, mycelial harvests, DNA extractions and sequencing runs (Supplementary Table S1). Based on the variation between parental replicates, individuals more than 95% similar to each other were considered clones, and one randomly selected individual from each clonal group was retained in the clone-corrected data set.

### Population Structure

Principal component analysis was performed on a scaled and centered genotype matrix in the R package pcaMethods (Stacklies et al., 2007), using the *nipalsPCA* method to account for the small amount of missing data (method=’nipals’, center=TRUE, scale=’uv’). This method was used for all PCAs performed. To estimate pairwise differentiation between years, we used Weir and Cockerham’s (1984) F_ST_ measure (Weir and Cockerham, 1984), which weights allele frequency and variance estimates by population size, implemented in the R package *StAMMP* with the *stamppFST* function (Pembleton et al., 2013). We performed 1000 permutations of the SNP set to assess if F_ST_ estimates were significantly greater than zero (Pembleton et al., 2013; Weir and Cockerham, 1984).

### Measures of inbreeding

We used the canonical method-of-moments estimator of the individual inbreeding coefficient, F=l-H_o_/H_e_, where H_o_ is the observed individual heterozygosity, and H_e_ is the expected heterozygosity given allele frequencies in a reference population assumed to be at HWE (Keller et al., 2011; Purcell et al., 2007). We utilized allele frequencies in the *in vitro* F_1_ to define expected heterozygosity, a theoretical equivalent to evaluating F with respect to the parental allele frequencies (Wang, 2014). For each isolate, F was calculated with respect to non-missing genotypes only. To compare average F between years, a pairwise t-test was implemented in R with *pairwise.t.test* (pool.sd=FALSE, paired=FALSE, p.adjust.method=’bonferroni’).

Heterozygosity was defined as the number of isolates with a heterozygous genotype at each SNP divided by the total number of non-missing genotype calls. Minor allele frequency (MAF) was defined as the number of minor alleles, where the minor allele was defined as the allele with the lowest frequency in the field population, present at each SNP divided by the total number of non-missing chromosomes (number of non-missing genotype calls multiplied by two). Heterozygosity and MAF distributions for each year and the *in vitro* F_1_ were graphically assessed using the *density* function in R.

For each individual, we calculated the proportion of MEs, defined as the ratio of MEs to the total number of non-missing tested sites, analogous to the PLINK implementation (–mendel) (Purcell et al., 2007). An ME was defined as a genotype inconsistent with the individual being an F_1_ derived from the two founding parental isolates (Purcell et al., 2007). An isolate with a proportion MEs exceeding the *in vitro* F_1_ mean by 3 s.d. was classified as field inbred and otherwise as field F_1_.

### Genome scan for allele frequency differentiation

To detect regions of differentiation between the *in vitro* F_1_, field F_1_, and inbred isolates, we performed a Fisher’s exact test of allele counts for all pairwise comparisons, using the *fisher.test* function in R. *P*-values were adjusted for multiple testing using the Benjamini and Hochberg (1995) procedure, implemented with the *p.adjust* function, at a false discovery rate (FDR) of 10% (Benjamini and Hochberg, 1995; Wright, 1992). Significant SNPs were retained in further analyses only if another SNP within 200 kb also surpassed the significance threshold.

We compared the F_ST_ distribution of significantly differentiated SNPs within ROIs to the genome-wide F_ST_ distribution according to Lewontin and Krakauer (1973). Here, F_ST_ wasdefined as, 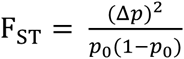, where *p*_*0*_ is the frequency of the minor allele in the field F_1_, and *Δp* is the difference in allele frequency between the field F_1_ and field inbred subpopulations.

Haploview (Barrett et al., 2005) was used to estimate pairwise LD (*r*^*2*^) between SNPs in scaffolds containing ROIs.

### Haplotyping

As the population was established by two parental strains, assuming no mutation, all isolates were by definition combinations of the founding parental haplotypes. Therefore, we took a deterministic approach to phasing, akin to utilizing trio information to phase parental genotypes (Browning and Browning, 2011). Haplotyping in regions of interest was further facilitated by the fact that either or both parents were homozygous, with the homozygous genotype assumed to represent a founding parental haplotype. We showed that this was a valid assumption by analyzing early replicates of the parental genotypes that represented the “ancestral” heterozygous genotype in a specific region (Supplementary Figure S13) and by comparison to homozygous genotypes of selfed isolates (data not shown). We used the homozygous parental stretches (haplotypes) to deduce the other haplotypes from consensus genotypes for the expected genotypic classes. Progeny membership in a genotypic class was defined by *k*-means clustering using the *kmeans* function in R (centers=8, n.iter=1000, nstart=100). To further refine clusters and remove recombinant isolates, we calculated local pairwise relatedness, defined as IBS, between isolates within a cluster, and removed isolates that shared on average less than 90% IBS with the respective cluster members. Next, we defined the consensus genotype based on the refined clusters utilizing the majority rule (see “Identifying a clone-correction threshold”), and heterozygous genotypes within haplotypes were set to missing.

To determine the haplotype composition of each isolate, the three identified haplotypes in a region of interest were used to construct reference genotypes for all possible haplotype combinations (e.g. H1/H2, H1/H1). Then, the genotypic discordance (i.e. the number of mismatched genotypes) between each isolate genotype and reference genotype were calculated. The most similar reference genotype was assigned if genotypic discordance was less than 25%. Otherwise, the isolate genotype was deemed “Unknown.”

To create phase diagrams, haplotype tagging SNPs (SNPs which unambiguously distinguished a specific haplotype) were identified at SNPs where all haplotypes had no missing data. Individual genotypes were then classified for homozygosity or heterozygosity at each haplotype tagging SNP.

### Identifying mating type associated SNPs

We performed a Fisher’s exact test of allele frequency differences between isolates of opposite mating types in the field F_1_. Multiple test correction was performed as above (see ‘Genome scan for allele frequency differentiation’).

### Heterozygosity in the MTR

To test differences between the heterozygote frequency distribution in the mating type region relative to the rest of the genome, we compared the heterozygosity of genome-wide SNPs sampled in equal proportions of marker types (e.g. *AA* x *Aa*) to the mating type region using the *sample* function in R without replacement (replace=FALSE), excluding SNPs not polymorphic or with missing data in the parental isolates. The identified ME-enriched SNPs were excluded (see Supplementary Text). To account for an unequal ratio of A1 to A2 mating type isolates, in each test, the A2s were down-sampled (without replacement) to equate with the A1 sample size in the respective subpopulation. We used the *wilcox.test* function in R to perform a one-sided, unpaired Wilcoxon rank sum test (alternative=’less’, paired=FALSE), repeated for 100 random SNP samples. Additionally, heterozygosity distributions of the A1 and A2 isolates in each subpopulation were compared to the respective genome-wide distribution, with SNP but not isolate down-sampling.

Heterozygote excess was tested at each locus in the A1 and A2 isolates for each subpopulation, using the function *HWExact* in the R package HardyWeinberg (Graffelman, 2015; Wigginton et al., 2005). This amounts to a one-sided test of HWE where heterozygote excess is the only evidence of deviation from HWE. We controlled for multiple testing as above.

### Allele frequency changes in the MTR

In the MTR, the frequency of the parental tagging allele (*p*_*a*_)at SNPs heterozygous in one parent and homozygous in the other, was calculated for A1 and A2 isolates separately in the parental generation, the field F_1_ and the field inbreds, excluding missing genotypes. The A2 tagging SNPs were separated into two categories based on *p*_*a*_ with respect to mating type in the field F_1_. The First case consisted of SNPs with *p*_*a*_≥0.3 in the A2 isolates and *p*_*a*_≤0.3 in the A1 isolates, and the second case consisted of the remaining SNPs. Expectations for *p*_*a*_ in theoretical F_1_ and F_2_ populations, for the three cases where the *a* alleles is in the haplotype background of the: 1) Y in the male sex; 2) X in the male sex; and 3) X in the female sex, were derived based on the formulas in (Allendorf et al., 1994; Clark, 1988).

## Acknowledgements

We thank Daniel C. Hut and Amara R. Dunn for helpful conversations. We thank Holly W. Lange and Michael R. Fulcher for assistance with field and lab work. This work was supported by the New York State Department of Agriculture and Markets (grant numbers C200780, C200818 to C.D.S.). This work was also partly funded by USDA-NIFA/DOE Biomass Research and Development Initiative (BRDI) (grant number 2011-06476 to M.A.G.) and Cornell University startup funds to M.A.G.

### Author Contributions

CDS and MOC conceived the experimental design; MOC, CDS, EG, and MAG conceptualized the analysis; MOC performed the experiments and analyses; MOC and EG wrote the manuscript; CDS and MAG revised the manuscript.

**Supplementary Materials**

**Supplementary Text. Additional methods and results.**

**Supplementary Tables S1-S8.**

**Supplementary Figures S1-S18.**

